# Schizophrenia-like neurodevelopmental pathology reshapes experience-dependent brain network remodeling following adolescent alcohol exposure

**DOI:** 10.64898/2026.08.26.747060

**Authors:** Charles Houdant, Maedeh Khalilian, Zoé Fortineau, Cassandre Rouanet, Elise Leuillier, Margaux Madeline, Sidy Fall, Ardalan Aarabi, Jérôme Jeanblanc, Mickaël Naassila

## Abstract

**Background:** Alcohol use disorder (AUD) is highly prevalent in schizophrenia, yet the neurobiological basis of this vulnerability remains poorly understood. Neurodevelopmental models suggest that pre-existing brain dysconnectivity may increase vulnerability to AUD. We therefore tested whether schizophrenia-like neurodevelopmental pathology alters how alcohol-related experience is incorporated into large-scale brain networks.

**Methods:** Resting-state functional connectivity was assessed in male Sprague-Dawley rats (*n* = 18-21/group) with neonatal ventral hippocampal lesions (NVHL), a neurodevelopmental model of schizophrenia, and sham-operated controls, with or without voluntary adolescent alcohol exposure. Functional connectivity was assessed using seed-to-voxel and seed-to-seed analyses within a cortico-striato-limbic network. We additionally examined whether individual alcohol intake during adolescence predicted adult functional connectivity according to neurodevelopmental status.

**Results:** NVHL and adolescent alcohol exposure independently produced predominantly hypoconnected cortico-striato-limbic networks. However, alcohol exposure did not exacerbate NVHL-associated dysconnectivity but was instead associated with a distinct network organization characterized by relative hyperconnectivity. Although alcohol intake was comparable between groups, dose-dependent relationships between adolescent alcohol consumption and adult functional connectivity were observed in sham animals but were absent or markedly attenuated in NVHL rats. These effects were primarily centered on prelimbic cortex connectivity with the amygdala, hippocampus, and dorsal striatum, implicating this circuitry in altered experience-dependent remodeling.

**Conclusions:** These findings suggest that vulnerability to AUD associated with schizophrenia-like neurodevelopment may arise less from additive network dysfunction than from an altered capacity of large-scale brain networks for experience-dependent functional remodeling. Schizophrenia-like neurodevelopmental pathology may therefore change how alcohol-related experience is translated into persistent brain network organization.

## 1. Introduction

Alcohol use disorder (AUD) is among the most frequent and detrimental comorbidities of schizophrenia, occurring two to three times more often than in the general population and contributing to poorer functional outcomes, reduced treatment adherence, and increased mortality (1–3). Despite this burden, the neurobiological mechanisms underlying this vulnerability remain poorly understood. Current neurodevelopmental models propose that schizophrenia-related alterations of cortico-striato-limbic circuits increase susceptibility to addictive disorders by altering reward processing and executive control (4,5). An alternative possibility is that neurodevelopmental pathology fundamentally alters how alcohol-related experience is translated into long-term brain network organization.

To investigate this comorbidity, the neonatal excitotoxic lesion of the ventral hippocampus (NVHL) rat model provides a relevant neurodevelopmental framework. Translational evidence suggests that excessive ventral hippocampal glutamatergic activity may drive the emergence and progression of psychosis (6). Consistent with this framework, NVHL rats exhibit delayed post-adolescent psychosis-like abnormalities, negative-like and cognitive deficits across development, and widespread mesocorticolimbic dysfunction (7). NVHL rats also show enhanced vulnerability to addictive drugs, including cocaine (8–10) and nicotine (11). Importantly, this model enables dissociation between the amount of alcohol exposure and how a neurodevelopmentally altered brain incorporates that experience into its subsequent functional organization (12).

Resting-state functional MRI (rs-fMRI) offers a systems-level framework for investigating how distributed brain circuits are organized following developmental and environmental perturbations. Beyond identifying dysconnectivity, resting-state connectivity captures large-scale functional architecture reflecting experience-dependent network plasticity and developmental refinement (13). In both schizophrenia and AUD, converging evidence indicates widespread disruption of cortico-striato-limbic networks supporting executive control, contextual processing and reward-related behaviors (14–19). To date, however, only one MRI study has examined schizophrenia–AUD comorbidity, reporting persistent glutamatergic and GABAergic abnormalities in the cingulate cortex and nucleus accumbens (NAc) in the NVHL model (20). Whether schizophrenia-like neurodevelopment fundamentally modifies the long-term network consequences of adolescent alcohol exposure nevertheless remains unknown.

Accordingly, we combined rs-fMRI with the NVHL model of schizophrenia–AUD comorbidity to investigate whether neurodevelopmental pathology alters the functional architecture of cortico-striato-limbic circuits and how adolescent alcohol exposure becomes embedded within these networks. Using complementary seed-to-voxel and seed-to-seed analyses, we characterized the independent and combined effects of NVHL and adolescent alcohol exposure on cortico-striato-limbic functional organization. We then tested the central prediction arising from this framework: namely, whether the relationship between the amount of alcohol consumed during adolescence and subsequent functional connectivity differs according to neurodevelopmental status. Specifically, if schizophrenia-related vulnerability reflects impaired experience-dependent network remodeling rather than additive dysconnectivity, the amount of alcohol consumed during adolescence should predict adult functional network organization in control animals but not following neurodevelopmental hippocampal lesions.

## 2. Methods and Materials

### 2.1. Animals

Male Sprague-Dawley rats were bred from 16 pregnant dams obtained from Charles River (Saint-Germain-Nuelles, France). Eighty-six male pups from two independent cohorts were housed individually with environmental enrichment (cardboard tunnel and wood stick, music on display in the facility) after weaning on postnatal day (PND) 21, under controlled temperature (22 ± 1°C) and humidity conditions, with food and water available *ad libitum*. Animal welfare procedures are detailed in the Supplementary Methods. Procedures complied with European Union Directive 2010/63/EU, were approved by the local ethics committee (CREMEAP; APAFIS#34211), and followed the ARRIVE guidelines.

### 2.2. Surgery

On PND 7, pups underwent bilateral NVHL or SHAM surgery in a stereotaxic apparatus (Kopf Instruments, USA) under isoflurane anesthesia, as previously described (12,20). NVHL pups (*n* = 44) received ibotenic acid into the ventral hippocampus (Hipp) (10 mg.mL⁻¹ in artificial cerebrospinal fluid, 0.3 µL per side, 0.15 µL.min^-1^, Tocris Bioscience, UK), whereas SHAM pups (*n* = 42) received artificial cerebrospinal fluid (Harvard Apparatus) at the same coordinates (AP - 3.0 mm, ML ±3.5 mm, DV-5.0 mm from bregma). Surgical details are provided in the Supplementary Methods. Two pups died during surgery.

### 2.3. Adolescent alcohol exposure

Adolescent alcohol exposure was performed as previously described (12). From PND 28 to PND 42, alcohol-exposed rats had continuous 24-hour access to two bottles, one containing 10% ethanol (v/v) and the other tap water, whereas control rats had access to tap water only (see Supplementary Methods).

### 2.4. Rs-fMRI

#### 2.4.1 Image acquisition

Rs-fMRI was performed between PND 55 and PND 72, corresponding to the late-adolescent/early-adult period when schizophrenia-related phenotypes emerge in the NVHL model (7). Rats were anesthetized with isoflurane in air (1 L.min^-1^, 5% induction, 1.5–1.8% maintenance), with continuous monitoring of heart rate, respiration, and temperature. Anesthesia was adjusted to maintain stable respiration. Images were acquired on a 7 T BioSpec 70/20 USR system (Bruker, Ettlingen, Germany). Coronal functional images were obtained using a spin-echo echo-planar imaging (EPI) sequence (TR/TE = 2000/16 ms, flip angle = 90°, 450 repetitions, FOV = 30 × 25 mm², matrix = 64 × 64, 34 slices, in-plane spatial resolution = 0.48 × 0.40 mm², 0.8 mm slice thickness, and no interslice gap), followed by a high-resolution T2-weighted (T2w) anatomical scan. Additional acquisition and monitoring details are provided in Supplementary Methods.

#### 2.4.2. Segmentation and volumetric analysis

Manual segmentation was performed on T2w anatomical images using 3D Slicer software (v5.8.0; https://www.slicer.org/). Regions of interest (ROI) included the whole-brain reference mask, the total Hipp, and the left and right Hipp (see Supplementary Methods). Relative hippocampal volume was calculated by normalizing hippocampal volume to whole-brain volume. In NVHL rats, left and right hippocampal volume loss was expressed as the percentage reduction relative to the corresponding hemisphere in SHAM rats.

#### 2.4.3. Image preprocessing

Data were preprocessed using the Rodent Whole-Brain fMRI Data Preprocessing Toolbox (21), following previously described procedures (22) integrating AFNI (https://afni.nimh.nih.gov/), FSL (v5.0, https://www.fmrib.ox.ac.uk/fsl), and ANTs (http://stnava.github.io/ANTs/). Preprocessing included image preparation, slice-timing and motion correction, bias-field correction, brain extraction, nuisance regression, temporal filtering (0.01–0.1 Hz), spatial normalization, and smoothing. The mean EPI image was registered directly to the SIGMA-Wistar rat EPI template (23). Nuisance regression was performed in native functional space before normalization, with template-derived tissue masks transformed into native space for nuisance-signal extraction. The estimated transforms were subsequently applied to the filtered 4D data, followed by spatial smoothing with a 3-mm FWHM Gaussian kernel. Registration quality and tissue-mask placement were visually inspected for all animals. Full preprocessing and quality-control details are provided in the Supplementary Methods and Figure S1.

#### 2.4.4. Image postprocessing

Resting-state functional connectivity was assessed using complementary seed-to-voxel and ROI-based seed-to-seed approaches (Figure 1). Seed-to-voxel analysis was used as an exploratory method to characterize the spatial distribution of functional connectivity from predefined seeds. ROI-to-ROI analyses provided a hypothesis-driven quantification of connectivity between these regions. Fourteen ROIs, comprising left and right prelimbic cortex (PL), infralimbic cortex (IL), NAc, dorsal striatum (DS), Hipp, amygdala (Amy) and basolateral Amy (BLA), were selected *a priori* based on involvement in AUD and schizophrenia (4,24,25). ROIs were defined according to the Duke Center for In Vivo Microscopy rat brain atlas (26). Further details on image postprocessing are provided in the Supplementary Methods.

**Figure 1.**
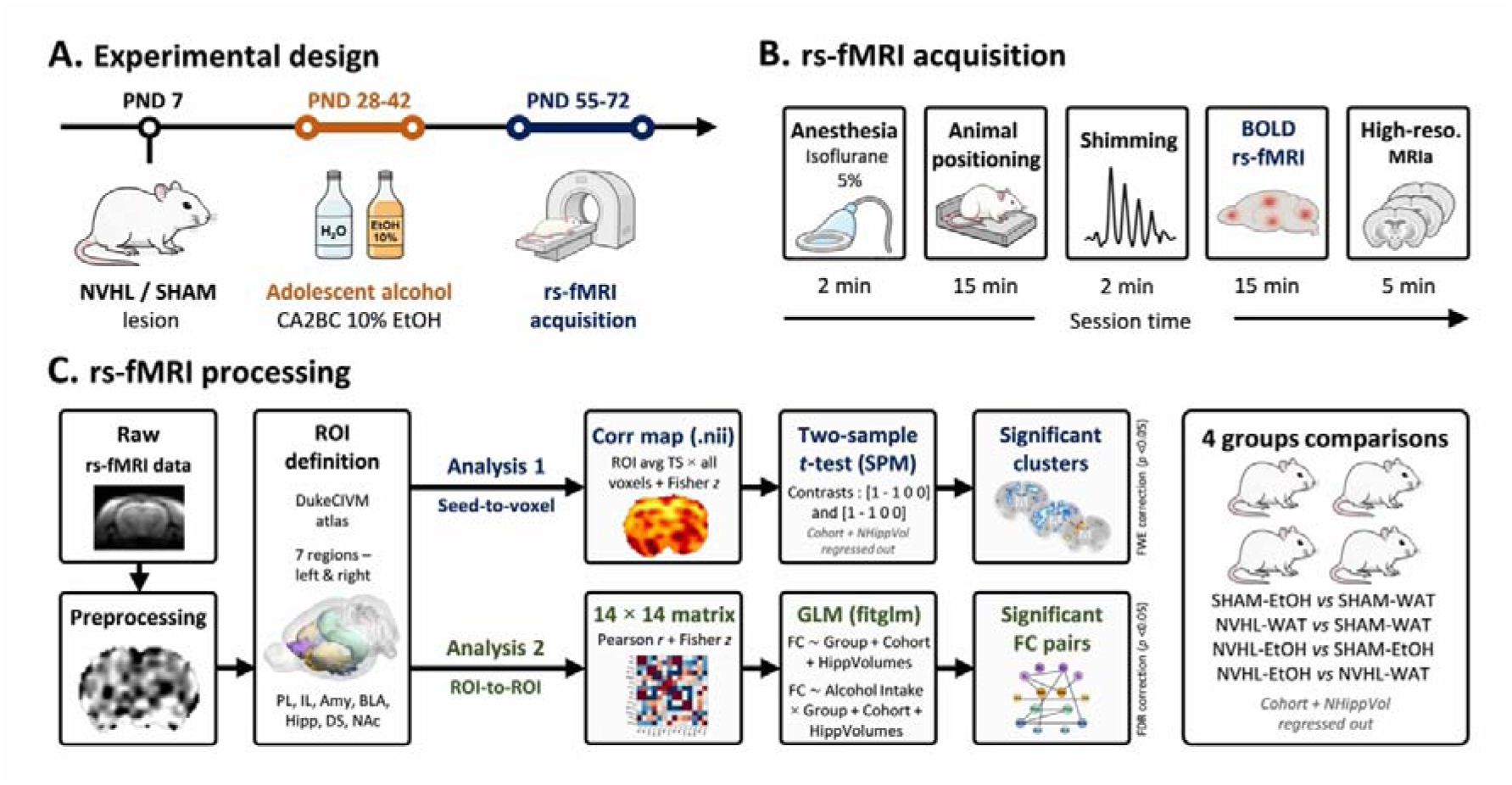
Experimental timeline and rs-fMRI analysis workflow. **(A)** Experimental design showing NVHL or SHAM surgery, adolescent alcohol exposure, and rs-fMRI acquisition. The continuous access two-bottle choice with 10% ethanol (CA2BC 10% EtOH) procedure initially generated four experimental groups: SHAM rats that were exposed to water (SHAM-WAT, *n* = 20) or ethanol (SHAM-EtOH, *n* = 21) during adolescence, and NVHL rats that were exposed to water (NVHL-WAT, *n* = 21) or ethanol (NVHL-EtOH, *n* = 22) during adolescence. **(B)** Rs-fMRI was acquired under isoflurane anesthesia with physiological monitoring, field shimming, BOLD rs-fMRI acquisition and T2w anatomical MRI (MRIa). Final analyzed sample sizes were SHAM-WAT *n* = 19, SHAM-EtOH *n* = 18, NVHL-WAT *n* = 18 and NVHL-EtOH *n* = 21. **(C)** Raw rs-fMRI data were preprocessed using a dedicated rodent whole-brain preprocessing workflow. After preprocessing, ROIs were defined and two complementary connectivity approaches were performed. Seed-to-voxel analyses generated whole-brain functional connectivity maps from each ROI time series, whereas ROI-to-ROI analyses quantified pairwise functional connectivity between the 14 predefined bilateral ROIs. Group-level analyses were then used to identify significant clusters and connectivity pairs across the four planned comparisons. Cohort and normalized hippocampal volume (NHippVol) were included as covariates in all group-level statistical analyses. Together, these analyses were used to characterize the effects of adolescent alcohol exposure, NVHL status, and their combination on resting-state functional connectivity (FC).

##### 2.4.4.1. Seed-to-voxel analysis

Whole-brain seed-to-voxel functional connectivity maps were generated for each ROI and animal by correlating the mean blood oxygenation level-dependent (BOLD) time series of the seed with voxelwise BOLD signals, followed by Fisher’s *r*-to-*z* transformation. Group analyses were performed in SPM (https://www.fil.ion.ucl.ac.uk/spm/) using two-sample *t*-tests, with cohort and normalized hippocampal volume entered as covariates. Each comparison was tested bidirectionally to identify higher or lower functional connectivity in the first group relative to the second. Four planned comparisons were tested: the effect of alcohol in SHAM rats (SHAM-EtOH *vs*. SHAM-WAT), the NVHL effect (NVHL-WAT *vs*. SHAM-WAT), the effect of alcohol between NVHL and SHAM rats (NVHL-EtOH *vs*. SHAM-EtOH), and the effect of alcohol in NVHL rats (NVHL-EtOH *vs*. NVHL-WAT). Significant clusters comprised contiguous voxels surviving a voxel-level family-wise error (FWE) correction threshold of *p*FWE < 0.05 and were anatomically localized using the rat brain atlas.

##### 2.4.4.2. Voxel-wise ROI-to-ROI analysis

ROI-to-ROI functional connectivity was estimated using a voxel-wise seed-to-target approach. For each ROI pair, voxel-wise Pearson correlations were Fisher *z*-transformed and averaged within the target ROI, and reciprocal seed-to-target estimates were averaged to obtain a symmetric connectivity value. Group differences were assessed using a general linear model (GLM) including cohort and normalized hippocampal volume as covariates for the four planned comparisons described above. Statistical significance was assessed using Freedman–Lane permutation inference (5,000 permutations), with false discovery rate (FDR) correction applied across the 91 ROI-to-ROI connections separately for each comparison (*p*FDR < 0.05).

##### 2.4.4.3. Group-dependent associations between adolescent alcohol consumption and functional connectivity

To assess associations between adolescent alcohol consumption and adult functional connectivity, pairwise connectivity among the 14 predefined ROIs was tested in GLMs including alcohol intake, group, their interaction, cohort, and normalized hippocampal volume. Only alcohol-exposed animals were included. Alcohol × Group interactions were assessed using Freedman–Lane permutation inference (5,000 permutations), with FDR correction across the 91 ROI-to-ROI connections (*p*FDR < 0.05). For exploratory characterization, connections with permutation *p* < 0.05 were further examined using group-specific *post hoc* regressions.

### 2.5. Postmortem histology

After the experimental procedures, animals were deeply anesthetized with 5% isoflurane and decapitated. Brains were removed, fixed in 4% paraformaldehyde, sectioned coronally through the Hipp and stained with Cresyl Violet to verify ventral hippocampal lesions in NVHL animals and absence of comparable damage in SHAM animals. Additional histological details are provided in Supplementary Methods.

### 2.6. Statistical analysis

Behavioral and structural MRI analyses were performed using SigmaPlot 11.0, and voxel-wise ROI-to-ROI connectivity analyses using MATLAB R2018b. Significance was set at *p* < 0.05. Normality and homogeneity of variance were assessed before analysis. Adolescent alcohol intake was analyzed using two-way repeated-measures ANOVA, relative hippocampal volume using two-way ANOVA, and hemispheric differences in lesion size using a paired *t*-test. Outliers were identified using Grubbs’ test with *α* = 0.05.

## 3. Results

### 3.1. No hemispheric difference in hippocampal volume loss in NVHL rats and comparable low adolescent alcohol intake between groups

Before examining functional connectivity, we first verified that adolescent alcohol intake was comparable between groups and characterized hippocampal volume loss following NVHL. Alcohol intake did not differ significantly between SHAM and NVHL rats (Figure 2A; F_(1,40)_ = 2.831, *p* = 0.100). Mean cumulative alcohol intake across five drinking sessions was 1.81 g.kg^-1^ in SHAM rats (range: 0.19–6.61 g.kg^-1^) and 2.67 g.kg^-1^ in NVHL rats (range: 0.23–7.37 g.kg^-1^). NVHL rats showed reduced relative hippocampal volume compared with SHAM rats (Figure 2B; F_(1,72)_ = 33.821, *p* < 0.001), with no effect of adolescent alcohol exposure (F_(1,72)_ = 0.684, *p* = 0.411) and no lesion status × adolescent alcohol exposure interaction (F_(1,72)_ = 0.576, *p* = 0.450). Hippocampal volume loss did not differ between hemispheres in NVHL rats (Figure 2C; t_(38)_ = - 1.380, *p* = 0.176).

**Figure 2.**
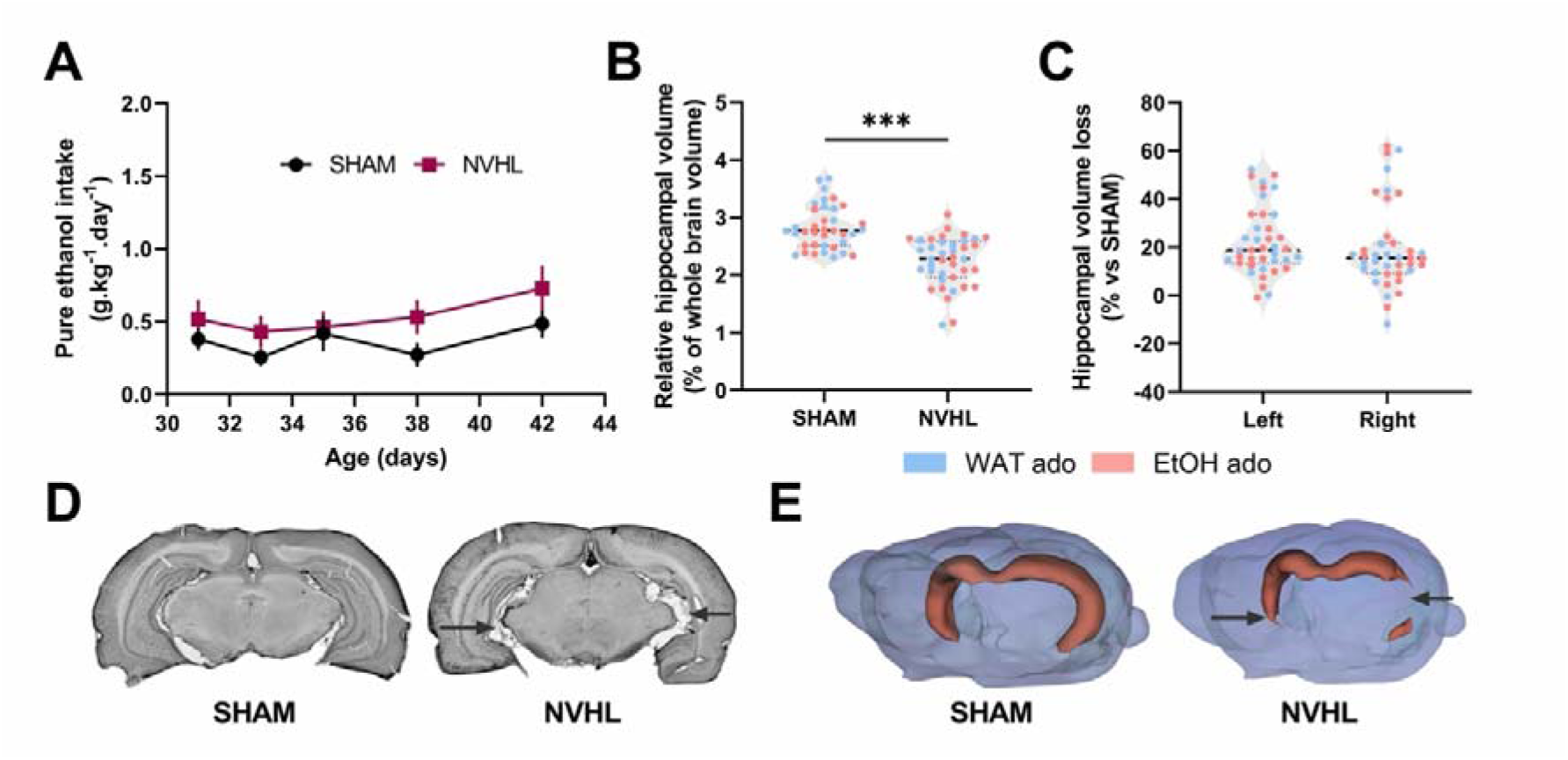
Adolescent alcohol intake, hippocampal volumetry, and anatomical verification of NVHL lesions. **(A)** Time course of alcohol intake during continuous voluntary access to 10% ethanol in SHAM and NVHL rats. **(B)** Relative hippocampal volume, expressed as a percentage of whole-brain volume, in SHAM and NVHL animals exposed to water (WAT) or ethanol (EtOH) during adolescence. **(C)** Left and right hippocampal volume loss in NVHL animals, expressed as a percentage relative to SHAM animals. **(D)** Representative Cresyl Violet-stained coronal sections from SHAM and NVHL animals, illustrating the absence of hippocampal damage in SHAM animals and bilateral ventral hippocampal lesions in NVHL animals. Arrows indicate lesion sites. (**E)** Representative 3D reconstructions of the whole-brain reference mask and Hipp in SHAM and NVHL animals, illustrating hippocampal tissue loss in NVHL animals. Data are shown as mean ± SEM or individual values with group mean, as appropriate. **\*\*\***p < 0.001.

### 3.2. NVHL and adolescent alcohol exposure produce reward-circuit hypoconnectivity but shift toward hyperconnectivity when combined

We next used seed-to-voxel analyses based on 14 predefined cortico-striato-limbic seeds (Figure 3A) to examine the independent and combined effects of adolescent alcohol exposure and neurodevelopmental pathology on large-scale functional connectivity. Although both factors separately produced qualitatively similar hypoconnectivity, their combination generated a markedly different network configuration (Figure 3B).

**Figure 3.**
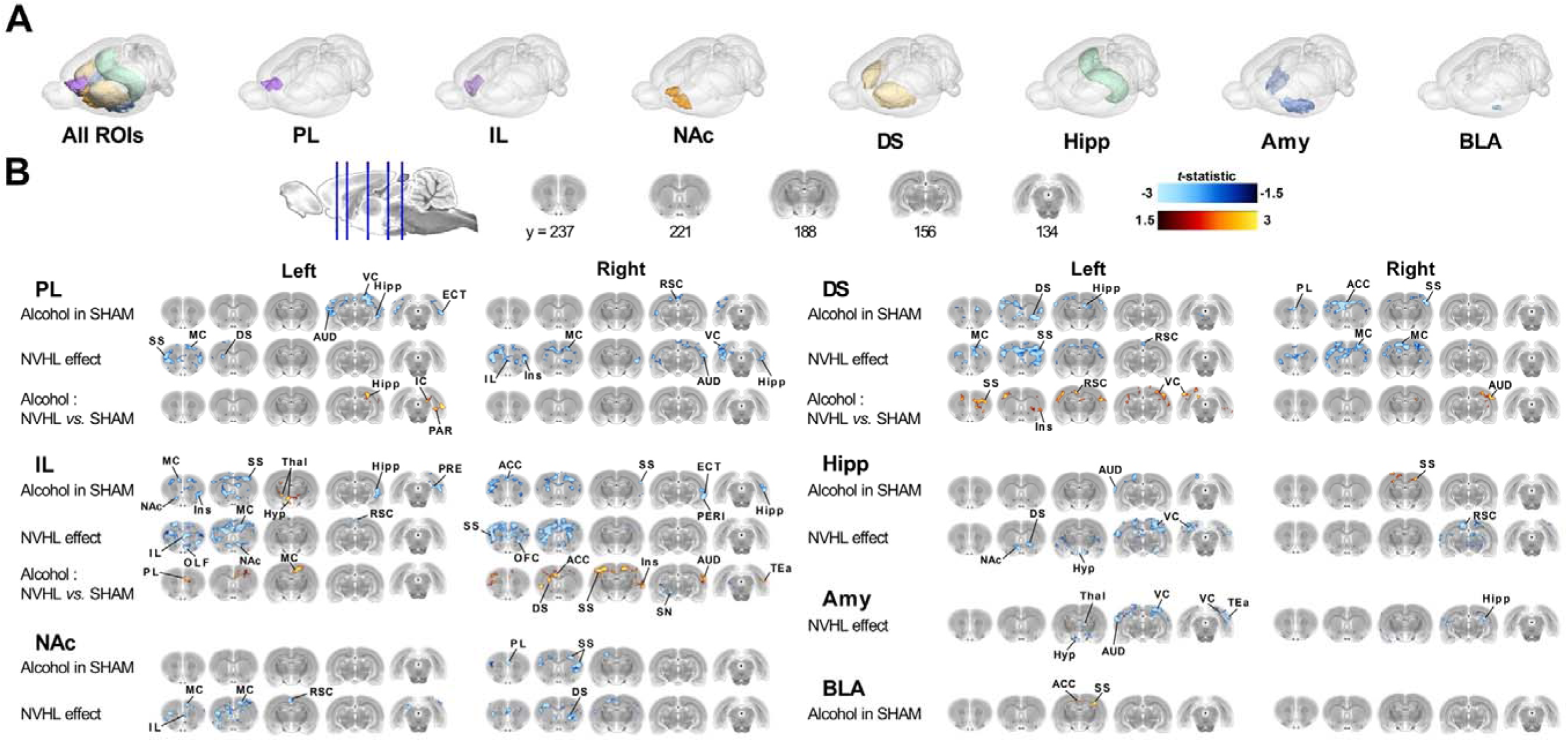
Seed-to-voxel mapping of altered resting-state functional connectivity from reward-circuit seeds. **(A)** Anatomical location of the reward circuit ROIs used as seeds for seed-to-voxel analyses, shown as 3D renderings. **(B)** Seed-to-voxel maps showing significant functional connectivity changes for each ROI across the indicated group comparisons. For clarity, only comparisons yielding significant clusters are displayed. Accordingly, the effect of alcohol in NVHL rats is not shown, as no significant clusters were detected for this comparison. Coronal slice levels are indicated at the top, with the displayed anatomical planes selected as representative of the whole-brain connectivity results. Left and Right indicate the hemisphere of the reference seed ROI used for the seed-to-voxel analysis. Statistical maps are color-coded according to the *t*-statistic, with red/yellow clusters indicating increased functional connectivity and blue clusters indicating decreased functional connectivity. ACC: anterior cingulate cortex; AUD: auditory cortex; ECT: ectorhinal cortex; Hyp: hypothalamus; IC: inferior colliculus; Ins: insular cortex; MC: motor cortex; OLF: olfactory cortex; PAR: parasubiculum; PERI: perirhinal cortex; PRE: presubiculum; RSC: retrosplenial cortex; SS: somatosensory cortex; TEa: temporal association cortex; Thal: thalamus; VC: visual cortex.

In SHAM animals, adolescent alcohol exposure predominantly reduced connectivity, mainly within prefrontal-striatal and intrastriatal circuits, particularly for IL and DS seeds. Similarly, the NVHL model produced predominant but more widespread hypoconnectivity involving prefrontal, striatal, hippocampal, and amygdalar regions. In both comparisons, significant reductions in connectivity also extended beyond the *a priori* selected reward-related regions to sensorimotor, sensory and associative cortical areas. Significant increases were sparse following alcohol exposure and even less frequent for the NVHL effect (whole-brain cluster-level results are provided in Table S1). By contrast, the comparison between alcohol-exposed groups revealed a predominantly hyperconnected profile in NVHL relative to SHAM animals, rather than an exacerbation of hypoconnectivity. Increased connectivity was mainly centered on IL and DS seeds and involved connections linking prefrontal regions, the DS, and the Hipp, with additional clusters extending to sensorimotor, sensory, and associative cortical areas. Only a limited number of significant negative voxels were detected, all associated with the right IL seed (Table S1). Finally, within the NVHL group, adolescent alcohol exposure produced no significant seed-to-voxel differences after voxel-level FWE correction, indicating attenuation of the widespread NVHL-related hypoconnectivity.

Seed-to-voxel alterations generally extended across both hemispheres, but bilateral concordance varied across comparisons and seeds (Table S1). The NVHL effect showed the most consistent bilateral organization, whereas the effect of alcohol in SHAM rats and, particularly, the comparison between alcohol-exposed NVHL and SHAM animals showed lower concordance between homologous regions and greater hemispheric specificity.

Voxel-wise ROI-to-ROI analyses were performed across the same 14 predefined regions used for the seed-to-voxel analyses (Figure 4A). Although no connection survived FDR correction, connectivity changes closely paralleled the whole-brain findings (Figure 4B; Table 1). Adolescent alcohol exposure in SHAM animals and the NVHL model of schizophrenia were both characterized by predominant ROI-to-ROI hypoconnectivity, with more widespread alterations in the NVHL condition. Likewise, comparing alcohol-exposed animals revealed an overall shift toward higher connectivity values in NVHL relative to SHAM rats, although the nominally significant connections included both increases and decreases. Finally, within the NVHL group, adolescent alcohol exposure largely attenuated the NVHL-related hypoconnectivity pattern, with only two connections reaching nominal significance.

**Figure 4.**
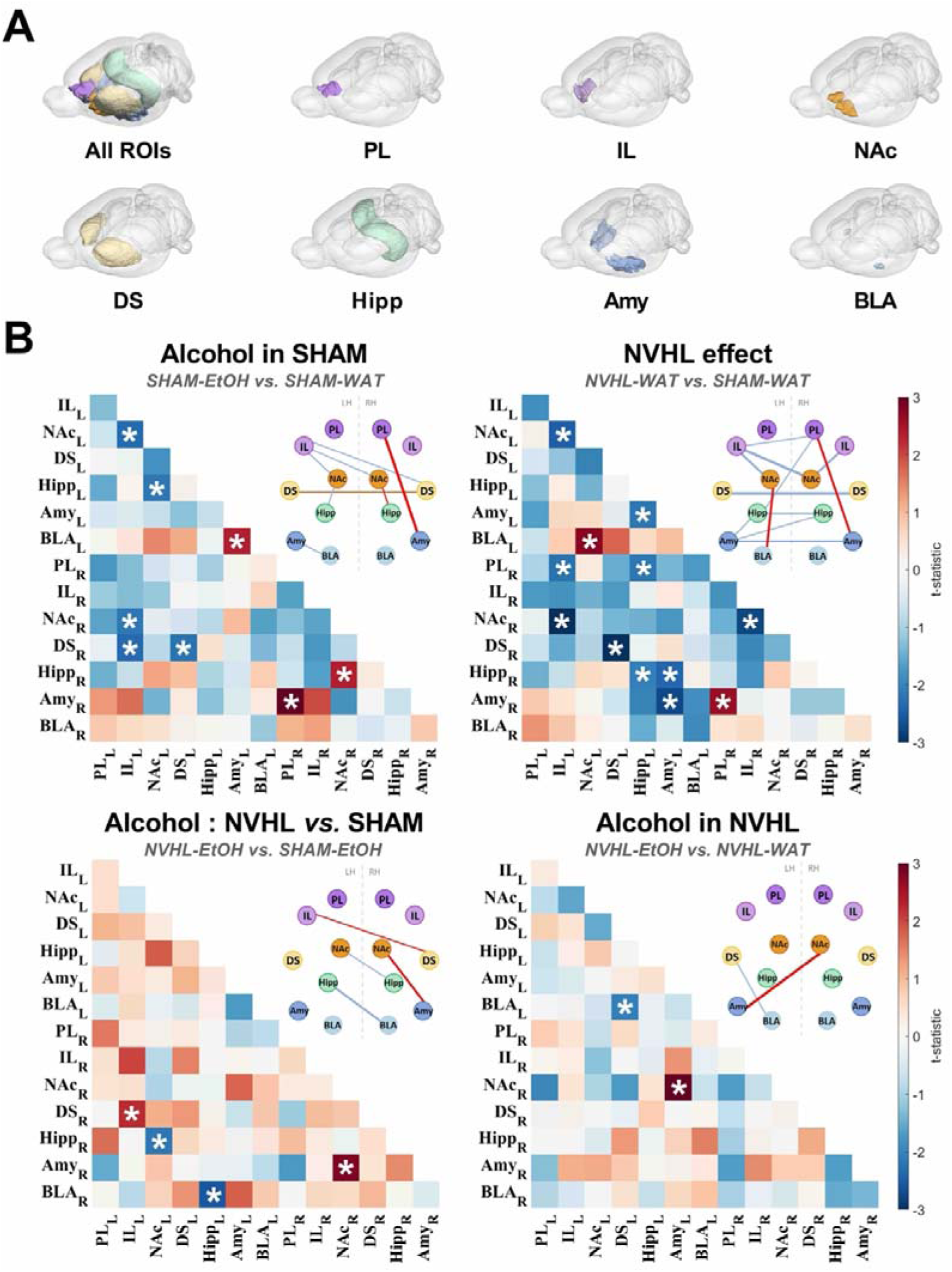
Voxel-wise ROI-to-ROI functional connectivity signatures of the reward-related circuit in alcohol exposure, schizophrenia, and comorbidity. **(A)** 3D visualization of the bilateral ROIs included in the functional connectivity analysis. **(B)** Matrices showing ROI-to-ROI functional connectivity differences for the four main comparisons. Values are expressed as *t*-statistics, with warm and cold colors indicating increased and decreased connectivity, respectively, in the first group relative to the second group. Asterisks denote connections with nominal significance based on permutation inference (uncorrected *p*_perm_ < 0.05). Network plots summarize the significant connections identified in each comparison. LH: left hemisphere, RH: right hemisphere.

**Table 1.** Statistical results of the voxel-wise ROI-to-ROI functional connectivity analyses shown in. **Figure 4**. Only connections with uncorrected *p*_perm_ < 0.05 are reported. For each connection, the table indicates the corresponding t-value, uncorrected *p*-value, Hedges’ *g* with 95% confidence interval, and direction of connectivity change.

| Comparison | ROI pair | *t*-value | $p_{perm}$<br>unc. | Hedges' *g* [95 % CI] | Connectivity<br>change |
| --- | --- | --- | --- | --- | --- |
| SHAM-EtOH<br>vs<br>SHAM-WAT | IL_L – NAc_L | -2.342 | 0.026 | -0.76 [-1.42, -0.09] | Decrease in<br>SHAM-EtOH |
|  | IL_L – NAc_R | -2.137 | 0.042 | -0.70 [-1.35, -0.03] |  |
|  | IL_L – DS_R | -2.344 | 0.015 | -0.76 [-1.42, -0.09] |  |
|  | NAc_L – Hipp_L | -2.183 | 0.033 | -0.71 [-1.37, -0.04] |  |
|  | DS_L – DS_R | -2.247 | 0.030 | -0.73 [-1.39, -0.06] |  |
|  | Amy_L – BLA_L | 2.365 | 0.023 | 0.77 [0.1, 1.43] | Increase in<br>SHAM-EtOH |
|  | PL_R – Amy_R | 3.171 | 0.003 | 1.03 [0.34, 1.71] |  |
|  | NAc_R – Hipp_R | 2.342 | 0.022 | 0.76 [0.09, 1.42] |  |
| NVHL-WAT<br>vs<br>SHAM-WAT | IL_L – NAc_L | -2.469 | 0.019 | -0.80 [-1.47, -0.13] | Decrease in<br>NVHL-WAT |
|  | IL_L – PL_R | -2.226 | 0.033 | -0.72 [-1.38, -0.06] |  |
|  | IL_L – NAc_R | -3.320 | 0.002 | -1.08 [-1.76, -0.38] |  |
|  | DS_L – DS_R | -3.081 | 0.003 | -1.00 [-1.68, -0.31] |  |
|  | Hipp_L – Amy_L | -2.161 | 0.041 | -0.70 [-1.36, -0.04] |  |
|  | Hipp_L – PL_R | -2.178 | 0.031 | -0.71 [-1.36, -0.04] |  |
|  | Hipp_L – Hipp_R | -2.087 | 0.045 | -0.68 [-1.33, -0.02] |  |
|  | Amy_L – Hipp_R | -2.406 | 0.022 | -0.78 [-1.44, -0.11] |  |
|  | Amy_L – Amy_R | -2.738 | 0.012 | -0.89 [-1.56, -0.21] |  |
|  | IL_R – NAc_R | -2.743 | 0.002 | -0.89 [-1.56, -0.21] |  |
|  | NAc_L – BLA_L | 2.765 | 0.010 | 0.90 [0.22, 1.57] | Increase in<br>NVHL-WAT |
|  | PL_R – Amy_R | 2.620 | 0.012 | 0.85 [0.18, 1.52] |  |
| NVHL-EtOH<br>vs<br>SHAM-EtOH | NAc_L – Hipp_R | -2.145 | 0.037 | -0.68 [-1.32, -0.03] | Decrease in<br>NVHL-EtOH |
|  | Hipp_L – BLA_R | -2.560 | 0.015 | -0.81 [-1.46, -0.16] |  |
|  | IL_L – DS_R | 2.223 | 0.030 | 0.71 [0.06, 1.35] | Increase in<br>NVHL-EtOH |
|  | NAc_R – Amy_R | 2.825 | 0.009 | 0.90 [0.23, 1.55] |  |
| NVHL-EtOH<br>vs<br>NVHL-WAT | DS_L – BLA_L | -2.049 | 0.046 | -0.65 [-1.29, 0] | Decrease in<br>NVHL-EtOH |
|  | Amy_L – NAc_R | 2.975 | 0.006 | 0.95 [0.28, 1.6] | Increase in<br>NVHL-EtOH |

### 3.3. NVHL alters the relationship between alcohol intake and subsequent functional network organization

The absence of group differences in adolescent alcohol intake enabled us to test our central hypothesis directly. We therefore examined whether interindividual variation in adolescent alcohol intake was differentially associated with adult functional connectivity in SHAM and NVHL rats. Although no Alcohol Intake × Group interaction survived FDR correction, ROI-to-ROI analyses identified several nominal interaction effects (*p_perm_* < 0.05) across several anatomically distinct connections (Figure 5A, C). Exploratory *post hoc* regressions demonstrated that these effects were primarily driven by alcohol intake–connectivity relationships in SHAM rats that were absent or markedly attenuated in NVHL rats (Figure 5A). In SHAM animals, higher alcohol intake was associated with stronger Amy–prefrontal and Hipp–NAc connectivity, but weaker PL–Hipp and PL–DS connectivity. By contrast, NVHL rats showed no comparable dose-dependent connectivity pattern. Together, these findings indicate that similar alcohol exposure becomes embedded within fundamentally different large-scale functional architectures according to neurodevelopmental status (Figure 5B).

**Figure 5.**
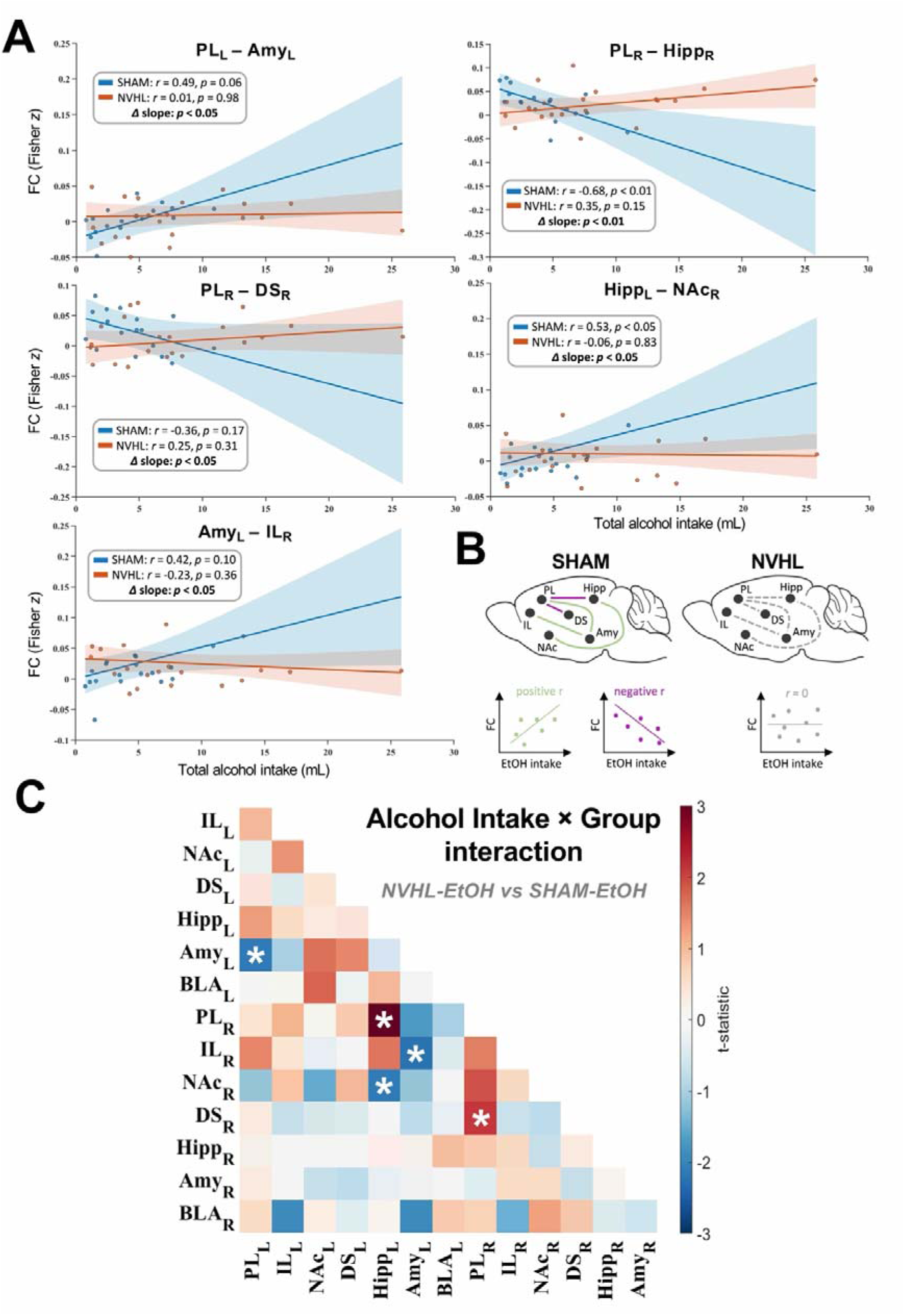
Adolescent alcohol intake–functional connectivity correlations in SHAM-EtOH and NVHL-EtOH rats. **(A)** Scatter plots illustrating the five ROI pairs showing significant Alcohol Intake × Group interactions. Each point represents one animal, with group-specific regression lines and 95% confidence intervals. Insets report group-specific partial correlation coefficients and associated *p* values, followed by the significance of the between-group difference in regression slopes. **(B)** Schematic representation of the group-specific alcohol intake–functional connectivity relationships. **(C)** Matrix showing Alcohol intake × Group interaction effects for ROI-to-ROI functional connectivity measures. Color intensity represents the *t*-statistic of the interaction term. Asterisks indicate connections showing nominal Alcohol Intake × Group interaction effects based on permutation inference (uncorrected *p*_perm_ < 0.05).

## 4. Discussion

Current neurodevelopmental models of schizophrenia–AUD comorbidity provide a compelling neurobiological framework but are largely built from converging evidence obtained separately in schizophrenia, AUD, and other substance use disorders. Consequently, the circuit-level mechanisms underlying this comorbidity have rarely been examined within a single experimental framework. Here, a fully crossed NVHL × adolescent alcohol exposure design revealed how schizophrenia-like neurodevelopment and alcohol exposure independently and jointly shape adult cortico-striato-limbic functional organization.

### 4.1. Independent NVHL-and alcohol-related hypoconnectivity in cortico-striato-limbic networks

Dissecting the brain network basis of schizophrenia–AUD comorbidity requires distinguishing the independent effects of NVHL and adolescent alcohol exposure. NVHL status produced a predominantly hypoconnected cortico-striato-limbic profile, robustly identified by seed-to-voxel analyses and qualitatively echoed by voxel-wise ROI-to-ROI results. This profile closely resembles resting-state findings in schizophrenia, which consistently report widespread reductions in functional integration across distributed networks (14,16). More broadly, it supports the dysconnectivity hypothesis, which conceptualizes schizophrenia not as isolated regional dysfunction but as impaired functional integration among interacting neural systems, whereby disruption of a key node may alter communication across broader networks (27,28). These findings extend the well-established behavioral, cognitive, and neurochemical phenotype of the NVHL model (7) by demonstrating persistent alterations in large-scale functional network organization, thereby further supporting its construct validity as a translational model of schizophrenia-related brain dysfunction (15,17,29).

Adolescent alcohol consumption was likewise associated with a predominantly hypoconnected profile, with the strongest effects for IL and DS seeds and reductions particularly involving prefrontal–striatal and intrastriatal connections. This organization is consistent with contemporary addiction models proposing that weakened prefrontal top-down inhibitory control, together with increasing DS recruitment, contributes to the transition from voluntary alcohol use to habitual and compulsive alcohol seeking (30). The extent of these alterations was particularly striking because SHAM animals consumed only 1.81 g.kg^-1^ of alcohol cumulatively over the two-week exposure period. Previous preclinical studies have similarly reported that alcohol exposure during adolescence produces long-lasting reductions in prefrontal–striatal connectivity involving the IL, PL, NAc, and DS (18,19). However, both studies used an intermittent binge-like regimen of 16 intragastric administrations of 5 g.kg^-1^ alcohol. Together, these observations suggest that even relatively low levels of alcohol exposure during adolescence may durably influence maturation of cortico-striato-limbic networks, particularly the prefrontal-striatal axis.

### 4.2. Schizophrenia-like neurodevelopmental pathology alters alcohol-related network reorganization and experience-dependent remodeling

The convergence of NVHL-and alcohol-related connectivity alterations within the cortico-striato-limbic network supports neurodevelopmental models of schizophrenia–AUD comorbidity (4,5). These models would predict that adolescent alcohol exposure further disrupts cortico-striato-limbic functional connectivity in NVHL rats. Consistent with the possibility of compounded neurobiological effects, structural MRI studies have reported more pronounced brain abnormalities in patients with schizophrenia and comorbid AUD than in those with either condition alone (31,32).

However, this prediction was not supported. In NVHL rats, adolescent alcohol exposure did not worsen hypoconnectivity. Rather, the widespread alterations observed in NVHL-WAT rats were no longer detected in NVHL-EtOH rats. This attenuation could initially suggest a compensatory effect of alcohol on pre-existing dysfunction. Individuals with schizophrenia may indeed report transient reductions in anxiety, dysphoria, or social discomfort after drinking (33,34). However, such reports reflect subjective relief rather than restoration of neurobiological deficits. Notably, controlled alcohol administration in patients with schizophrenia does not improve negative symptoms or cognitive performance and exacerbates some positive and perceptual disturbances (35). Our findings instead suggest a distinct pattern of relative hyperconnectivity within the neurodevelopmentally altered substrate. Indeed, NVHL-EtOH rats displayed predominant hyperconnectivity relative to SHAM-EtOH rats. Together, these findings argue against a simple additive interaction between schizophrenia-like neurodevelopment and alcohol exposure and instead support qualitative reorganization of functional networks determined by the pre-existing neurodevelopmental context.

The prominence of IL-and DS-centered alterations suggests that this reconfiguration may relate less to enhanced alcohol reward than to impaired behavioral regulation. The IL supports extinction and flexible updating of previously reinforced behavior (36,37), whereas progressive recruitment of the DS is thought to support automatic action selection as alcohol seeking becomes less sensitive to changes in outcome value or contingency (30). Such reorganization could weaken context-sensitive goal-directed control while favoring rigid, habit-like alcohol responding. This interpretation is consistent with evidence that latent inhibition and extinction abnormalities predict subsequent alcohol drinking in NVHL rats (38), and with our previous finding that comparable adolescent exposure produces an AUD-like phenotype in adulthood only in NVHL rats (12). At a complementary neurochemical level, the prominent prefrontal–striatal involvement observed here also complements previous spectroscopic findings in the same NVHL-based comorbidity model, which identified persistent GABAergic and glutamatergic abnormalities in the cingulate cortex and NAc after prolonged abstinence (20). Although rs-fMRI cannot establish causality, these findings identify a plausible circuit-level substrate through which schizophrenia-like neurodevelopment may facilitate later compulsive alcohol use.

A complementary picture emerged from the alcohol intake–functional connectivity analyses. Whereas group comparisons characterized mean connectivity differences between experimental conditions, these analyses examined whether subsequent network organization varied with the amount of alcohol consumed during adolescence and whether this relationship depended on neurodevelopmental status. In SHAM animals, alcohol intake scaled with subsequent connectivity across a specific subset of connections, suggesting that adolescent alcohol exposure left measurable signatures within the mature functional connectome. By contrast, these relationships were largely absent in NVHL animals despite comparable alcohol exposure. These findings suggest that neurodevelopmental pathology alters experience-dependent network remodeling rather than simply producing static functional dysconnectivity. Adaptive behaviors depend not only on the integrity of individual brain regions but also on the capacity of distributed neural systems to modify their interactions in response to experience and developmental refinement of large-scale functional organization (13,39,40). The marked attenuation of these nominal relationships in NVHL rats therefore suggests that neurodevelopmental abnormalities altered how alcohol-related experience was translated into subsequent network organization. This interpretation is consistent with models proposing that schizophrenia involves impaired synaptic plasticity in addition to large-scale dysconnectivity (41,42). Vulnerability to AUD may thus arise less from progressively increasing dysconnectivity than from reduced capacity of brain networks to undergo adaptive remodeling in response to alcohol-related experience.

Among the alcohol intake-dependent associations, the PL emerged as the principal hub, providing a potential circuit-level substrate for this interpretation. Through interactions with the DS, Hipp, and Amy, the PL integrates contextual and mnemonic information with the affective significance of experience, allowing previous experience to guide behavioral selection and updating (43). Such a function is particularly relevant to alcohol-related behavior, as the PL contributes to the context-dependent retrieval and expression of alcohol seeking (44). Loss of PL-centered intake–connectivity relationships in NVHL animals may therefore reflect a reduced capacity of these circuits to adapt proportionally to alcohol-related experience. Consequently, contextual, mnemonic and affective information may be less efficiently incorporated into behavioral updating. Consistent with our previous findings (12), such dysfunction could impair behavioral flexibility, favor the later persistence of alcohol-directed behavior, and contribute to the emergence of an AUD-like phenotype in NVHL animals.

Although most seed-to-voxel alterations involved bilateral networks, nominal ROI-to-ROI effects were often asymmetric and frequently involved interhemispheric connections. Because hippocampal volume loss was comparable between hemispheres, gross lesion asymmetry is unlikely to explain these observations. Instead, this pattern is compatible with the non-mirror-symmetric organization of rodent functional networks (45,46) and altered hemispheric functional organization reported in both schizophrenia and AUD (47–50). Although hemispheric specialization was not formally addressed here, these observations raise the possibility that altered hemispheric network organization contributes to schizophrenia–AUD comorbidity and deserves direct investigation.

### 4.3. Limitations and Future Directions

Although the present findings provide a circuit-level framework for schizophrenia–AUD comorbidity, several limitations should be considered. First, ROI-to-ROI analyses were restricted to an *a priori* cortico-striato-limbic network, and no individual connection survived correction for multiple comparisons. Nominal effects should therefore not be interpreted as definitive evidence for discrete abnormalities. Their convergence with FWE-corrected seed-to-voxel findings instead supports interpretation at the distributed-network level. Moreover, because this hypothesis-driven approach may overlook alterations outside the selected regions, future whole-brain analyses using graph-theoretical, connectome-wide, or independent component methods are needed to assess broader functional reorganization. Second, imaging at a single post-exposure time point prevented us from determining whether the observed alterations preceded, emerged during, or resulted from alcohol exposure. Repeated rs-fMRI before, during, and after adolescent alcohol exposure could clarify functional trajectories and identify early connectivity features associated with later alcohol-related behavior, using a longitudinal design similar to that previously applied with structural MRI in another neurodevelopmental model relevant to schizophrenia (51,52). Finally, because rs-fMRI provides an indirect measure of functional coupling (53), future studies combining effective connectivity approaches with causal circuit manipulations (e.g., optogenetics or chemogenetics) will be required to determine how the identified networks contribute to schizophrenia–AUD comorbidity.

### 4.4. Conclusion

To our knowledge, this study provides the first evidence that NVHL induces persistent alterations in cortico-striato-limbic functional connectivity and modifies the relationship between adolescent alcohol exposure and subsequent brain network organization. More broadly, these findings help refine neurodevelopmental models of schizophrenia–AUD comorbidity. Rather than representing a static vulnerability preceding alcohol exposure, schizophrenia-related dysconnectivity may determine how alcohol-related experience becomes incorporated into large-scale brain networks. Vulnerability to AUD may therefore arise jointly from atypical circuit organization and a reduced capacity of these networks to reorganize in response to experience. This framework may help explain why comparable alcohol exposure can lead to markedly different trajectories toward AUD. Importantly, our findings suggest that, within a neurodevelopmentally altered brain, the consequences of alcohol exposure may depend less on the amount consumed than on how that exposure is incorporated into brain network organization. These findings therefore emphasize the potential importance of limiting early alcohol exposure in individuals at risk for psychosis, before alcohol-related experience becomes incorporated into altered large-scale brain networks.

## Acknowledgments

CH has a PhD fellowship funded by the Université de Picardie Jules Verne in support to the FHU A^2^M^2^P. This work was supported by the FHU A^2^M^2^P, the INSERM and the CPER-MOSOPS (Contrat de plan État-région - Modélisation, Simulation, Optimisation des impacts, des Soins et des Parcours de Santé) funded by the Regional Council of the Hauts-de-France. We thank Virginie Jeanblanc for her valuable assistance with neonatal surgeries and the postoperative monitoring and recovery of the pups.

CH and MK: Methodology, Investigation, Formal analysis, Data curation, Visualization, Writing – original draft. ZF, CR, EL and MM: Investigation. SF: Investigation, Writing – original draft. AA: Conceptualization, Methodology, Data curation, Validation, Writing – review & editing, Supervision. JJ: Conceptualization, Methodology, Investigation, Data curation, Writing – original draft, Writing – review & editing, Supervision. MN: Conceptualization, Methodology, Data curation, Writing – original draft, Writing – review & editing, Supervision, Project administration, Funding acquisition.

The data supporting the findings of this study are available from the corresponding author upon reasonable request.

## SUPPLEMENTARY METHODS

### Animals and surgery

Animal welfare was monitored daily throughout the study using the Rat Grimace Scale. Following neonatal surgery, pups received intensified postoperative monitoring for five days using a clinical scoring system based on body weight, stomach filling, and behavior within the litter. Stomach filling was assessed by visual inspection of the abdominal milk spot, whose white appearance indicated adequate milk intake, while behavioral assessment included signs of abnormal isolation from littermates. Postoperative humane endpoints were determined according to this clinical score. On PND 7, pups weighing 12–20 g were anesthetized with isoflurane (Iso-Vet; 5% for induction and 2–3% for maintenance). Following a sagittal scalp incision, the skin was retracted to expose the skull. Pups were positioned in a stereotaxic apparatus equipped with an adaptor suitable for neonatal rats, and the head was stabilized using flat ear bars fitted with rubber protectors. Bilateral openings were drilled in the skull using a surgical drill (K.1070, Foredom, Germany) at the appropriate stereotaxic coordinates. Injectors (34-gauge, 25 mm length; Cooper’s Needle Works, UK) were then slowly lowered to the target depth. Following each infusion, injectors were left in place for 1 min to allow diffusion and minimize reflux before being slowly withdrawn. The surgical site was disinfected with an antiseptic solution (Vétédine, Vétoquinol, France), and the incision was closed using surgical glue (3M Vetbond, Phymep). Pups were maintained under a heat lamp until recovery from anesthesia and were then returned to their home cage.

### Adolescent alcohol exposure

During the adolescent exposure period, ethanol and water intake were monitored three times per week, on Mondays, Wednesdays, and Fridays, by weighing the bottles. Ethanol solution was prepared at 10% v/v in tap water using ethanol from VWR-Chemicals (Fontenay-sous-Bois, France). Body weight was recorded twice per week and daily average ethanol consumption was expressed in g·kg⁻¹·day⁻¹. To minimize potential side preference, the position of the ethanol and water bottles was alternated at each bottle-weighing session. Fluid loss unrelated to drinking was estimated using two additional cages without animals, each containing ethanol and water bottles, and consumption values were corrected for spillage. Of the six scheduled bottle-weighing sessions, one was excluded from all intake calculations because ethanol and water losses were abnormally elevated across the cohort and markedly exceeded the values observed during all other sessions, indicating a session-wide technical artifact unrelated to drinking. Cumulative adolescent ethanol intake was therefore calculated from the five remaining valid sessions. At the end of the exposure period, ethanol bottles were removed and animals were maintained under standard housing conditions until subsequent experiments. Animals were assigned to the four experimental groups according to a litter-balanced design to avoid confounding litter origin with lesion status or adolescent ethanol exposure. One SHAM-EtOH rat, whose average ethanol intake during adolescent exposure was approximately 16 times higher than the mean intake of the ethanol-exposed cohort and was identified as a significant outlier, was excluded from all subsequent analyses.

### Image acquisition

Animals were transferred to the MRI facility 15 min before imaging to allow acclimatization to the experimental environment. Animals were positioned prone on a Bruker rat bed assembly, and the head was secured using a bite bar and ear bars to minimize motion during image acquisition. Physiological parameters were monitored using an MR-compatible monitoring system (SA Instruments, Inc. Stony Brook, NY, USA). Body temperature was maintained at 35.8 ± 0.6°C using a warm-water circulation system (Bruker, Ettlingen, Germany). Heart rate (396 ± 31 bpm) and oxygen saturation were recorded using a fiber-optic pulse oximeter placed on the left hind paw. The isoflurane level was finely adjusted during acquisition to maintain a stable ventilation, with a mean respiratory rate of 81 ± 11 breaths.min^-1^. The 7 T BioSpec 70/20 USR system operated at 300 MHz with AV-III electronics and ParaVision360 v3.4. Radiofrequency transmission was performed using a 72-mm inner-diameter quadrature volume resonator, while signal reception was achieved with a four-channel phased-array receive-only surface coil (Bruker, Ettlingen, Germany). Magnetic field homogeneity was optimized using FASTMAP shimming. The Bruker standard EPI navigator was used with automatic ghost correction and drift compensation. Structural T2-weighted images were obtained using a fast spin-echo TurboRARE sequence, providing brain coverage with 28 contiguous coronal slices, each 0.8 mm thick. The other acquisition parameters were: RARE factor = 8; effective TR/TE = 2740/44 ms; FOV = 35 × 30 mm²; matrix size = 256 × 218; in-plane spatial resolution = 0.14 × 0.14 mm^2^; NEX = 4. The total acquisition time, including functional and anatomical scans, was approximately 23 min per animal. Datasets affected by poor image quality, acquisition artifacts, or technical problems were excluded from subsequent analyses. Rs-fMRI was performed between PND 55 and PND 72. The order of MRI acquisitions was counterbalanced across experimental groups to minimize age-related confounding. Mean age at scanning was 66.2 days in SHAM-WAT rats, 65.1 days in SHAM-EtOH rats, 63.9 days in NVHL-WAT rats, and 64.0 days in NVHL-EtOH rats.

### Segmentation and volumetric analysis

The segmentation procedure was adapted from previous manual volumetric approaches in rats, including studies conducted in control animals and in the NVHL model (1,2). All segmentations were performed by a blinded investigator with neuroanatomical expertise and reviewed by a senior investigator for anatomical consistency. Segmentations were performed on the native coronal images using visible anatomical landmarks and with reference to the rat brain atlas of Paxinos and Watson. The whole-volume reference mask was defined as the entire segmented tissue volume visible across the full rostrocaudal extent of the anatomical image series. Accordingly, all rostral and caudal sections included in the anatomical image series were retained, including the most rostral and caudal sections containing olfactory bulb and spinal cord tissue, respectively. Hippocampal segmentation included the hippocampal formation, comprising the cornu ammonis fields, dentate gyrus, and subiculum. Left and right hippocampi were segmented separately in order to assess hemispheric differences in hippocampal volume and lesion extent.

### Image preprocessing

The complete rs-fMRI preprocessing workflow is summarized in Figure S1. Data were preprocessed using the Rodent Whole-Brain fMRI Data Preprocessing Toolbox (https://github.com/GT-EmoryMINDlab/rodent-whole-brain-preprocessing-recipe). DICOM images were converted to NIfTI, reoriented, and spatial dimensions were rescaled by a factor of 10 to ensure compatibility with standard fMRI processing toolboxes (AFNI and FSL) (3,4). The first six volumes were discarded to allow signal stabilization. Slice-timing correction was performed with FSL slicetimer (TR = 2 s), followed by rigid-body motion correction using FSL MCFLIRT. A mean motion-corrected EPI image was generated for each animal, corrected for intensity inhomogeneity using N4 bias-field correction, and skull-stripped using FSL BET (fractional intensity threshold = 0.65). Brain masks were visually inspected and manually refined in FSLeyes. The masked mean EPI image was registered directly to the EPI template defined in SIGMA-Wistar rat brain space (5) using ANTs antsRegistrationSyNQuick.sh. Transforms were estimated before nuisance regression but were not applied to the full 4D BOLD series at this stage. Instead, inverse transforms were used to map template-derived cerebrospinal fluid (CSF) and white-matter (WM) masks into native functional space for nuisance-signal extraction. The resulting tissue masks were visually inspected on the individual mean EPI images and restricted to the individual brain masks. Native-space nuisance regression was performed using FSL fsl_glm. The nuisance model included constant, linear, and quadratic trends; six motion parameters estimated by MCFLIRT and their temporal derivatives; mean CSF and combined WM+CSF signals; and ten principal components extracted from non-brain tissue using AFNI 3dpc. To further account for non-brain fluctuations that were also expressed within the brain, non-brain principal components showing an association with the BOLD signal at *p* ≤ 0.001 in more than 1% of brain voxels were additionally included as nuisance regressors. Residual time series were scaled by the temporal mean and band-pass filtered between 0.01 and 0.1 Hz using AFNI 3dBandpass. The previously estimated ANTs transforms were then applied to the filtered residual time series using antsApplyTransforms, placing the data in common template space. Spatial smoothing was performed on the normalized, rescaled images using a Gaussian kernel of 3 mm FWHM. The normalized images were used for seed-based and ROI-to-ROI connectivity analyses. Cortical and subcortical regions of interest were defined using the Duke Center for In Vivo Microscopy rat brain atlas. The atlas was registered to the SIGMA-Wistar template space and used to extract regional time series for ROI-to-ROI functional-connectivity analyses. Registration quality was assessed by visual inspection of individual normalized mean EPI images overlaid on the rat EPI template and of inverse-warped CSF and WM masks in native functional space. Global brain boundaries and visible anatomical landmarks, including cortical contours, ventricular regions, hippocampal areas, and major subcortical structures, were inspected in all animals. Nonlinear deformation fields were additionally reviewed to identify implausible warping, with particular attention to hippocampal regions in NVHL animals. All data preprocessing and postprocessing were performed by the same investigator blinded to experimental group allocation.

### Voxel-wise ROI-to-ROI analysis

For each ordered ROI pair, the mean BOLD time series of the seed ROI was extracted and correlated with the BOLD time series of every voxel within the target ROI using Pearson correlation coefficients. The resulting coefficients were transformed into Fisher *z* values and averaged across all voxels of the target ROI, yielding one seed-to-target connectivity estimate for each ordered pair (ROI_i_as seed and ROI_j_as target). Because averaging voxel-wise correlations within the target ROI can produce asymmetric estimates depending on which ROI is treated as the seed, connectivity was calculated in both configurations, with ROI_i_as the seed and ROI_j_as the target, and vice versa. The final connectivity value for each ROI pair was defined as the arithmetic mean of these two reciprocal estimates, resulting in a symmetric ROI-to-ROI connectivity matrix. Analyses were restricted to the 14 predefined ROIs, yielding 91 unique ROI-to-ROI connections. Group differences in ROI-to-ROI connectivity were assessed using general linear models including group, cohort, and normalized hippocampal volume. The four prespecified group comparisons described in the main Methods were tested separately. Statistical significance was assessed using the Freedman–Lane residual-permutation procedure. For each connection and planned comparison, the full model included group, cohort, and normalized hippocampal volume, whereas the corresponding reduced model excluded the group effect while retaining cohort and normalized hippocampal volume. Residuals from the reduced model were randomly permuted across animals and added to its fitted values, after which the full model was refitted to each permuted dataset. This procedure was repeated 5,000 times to generate an empirical null distribution of the Group *t* statistic. Two-sided empirical permutation p values were calculated as (k + 1)/(Nperm + 1), where k was the number of permuted absolute *t* statistics greater than or equal to the absolute observed *t* statistic and Nperm = 5,000. Permutation-derived *p* values were subsequently corrected for multiple comparisons across the 91 unique ROI-to-ROI connections using false discovery rate (FDR) correction, separately for each planned group comparison, with statistical significance defined as *p*FDR < 0.05.

### Group-dependent associations between adolescent alcohol consumption and functional connectivity

Only alcohol-exposed animals were included in this analysis. For each ROI-to-ROI connectivity measure, functional connectivity was modeled as a function of adolescent alcohol intake, group, their interaction, cohort, and normalized hippocampal volume:

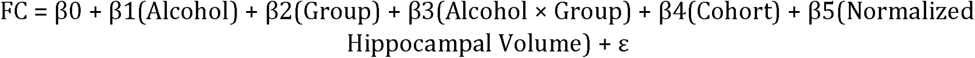

FC corresponded to the connectivity measure of interest, Alcohol to individual ethanol consumption during adolescence, and Group to lesion status among ethanol-exposed animals, coded as SHAM-EtOH or NVHL-EtOH. Cohort and normalized hippocampal volume were included to account for imaging cohort effects and variability in lesion extent. The Alcohol × Group interaction tested whether the relationship between adolescent alcohol consumption and functional connectivity differed between SHAM-EtOH and NVHL-EtOH animals. Statistical significance of the Alcohol × Group interaction was assessed using the Freedman–Lane residual-permutation procedure (6). For each connection, a reduced model excluding the interaction term, but retaining alcohol intake, group, cohort, and normalized hippocampal volume, was fitted. Residuals from this model were randomly permuted and added to the corresponding fitted values, after which the full model including the Alcohol × Group interaction was refitted. This procedure was repeated 5,000 times to generate an empirical null distribution of the interaction *t* statistic. A two-sided empirical permutation *p* value was calculated as (*k* + 1)/(*N*_perm_ + 1), where *k* was the number of permuted absolute *t* statistics greater than or equal to the absolute observed *t* statistic and *N*_perm_ = 5,000. Permutation-derived *p* values were subsequently corrected for multiple comparisons across the 91 ROI-to-ROI connections using FDR correction (*p*FDR < 0.05). For exploratory characterization of nominal interaction effects, connections with permutation *p* < 0.05 were further examined using separate within-group regressions to determine the direction of the association between alcohol consumption and functional connectivity in SHAM-EtOH and NVHL-EtOH animals.

### Postmortem histology

Following fixation, coronal sections containing the hippocampus were cut at a thickness of 100 µm using a vibratome (VT 1200S, Leica Microsystems, Nussloch, Germany). Sections were then stained with Cresyl Violet (Sigma-Aldrich) for cytoarchitectural visualization of hippocampal structures. Briefly, sections were rinsed in distilled water, incubated for 10 min at 37°C in Cresyl Violet solution, and rinsed again. Sections were then dehydrated through graded ethanol baths, cleared in xylene, and coverslipped using Entellan™ mounting medium (Merck/Sigma-Aldrich, Darmstadt, Germany; ref. 1.07960.0500). Representative images of mounted sections were acquired using a THUNDER Imager Tissue system (DM6-B, Leica Microsystems, Wetzlar, Germany) in brightfield/transmitted-light mode, using an N PLAN 5×/0.12 dry objective.

## SUPPLEMENTARY FIGURE

**Figure S1.**
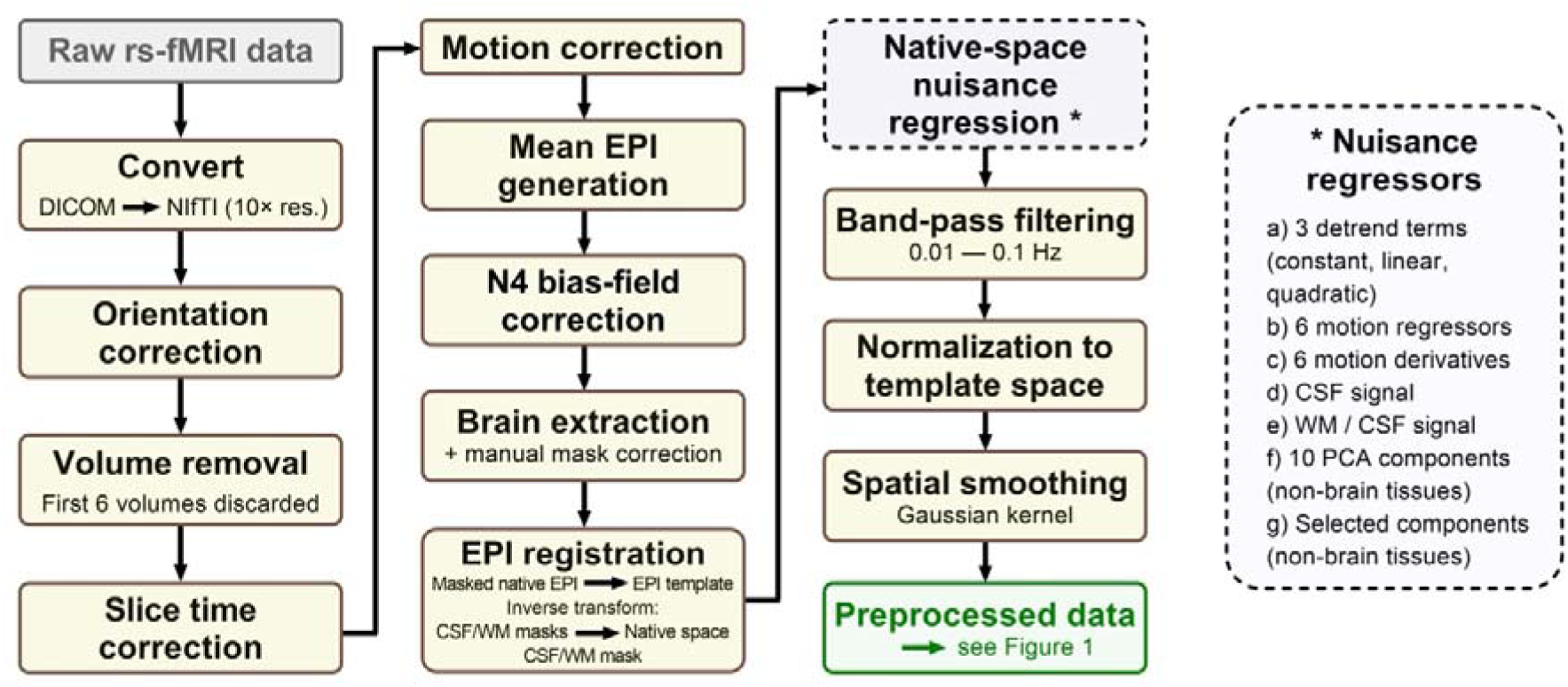
Rs-fMRI preprocessing workflow. Overview of the preprocessing pipeline applied to resting-state fMRI datasets before functional-connectivity analyses. Following image preparation, motion correction, bias-field correction, and brain extraction, the masked mean echo-planar imaging (EPI) image was registered to the SIGMA-Wistar EPI template. The estimated inverse transforms were used to map template-derived cerebrospinal fluid (CSF) and white matter (WM) masks into native functional space, where nuisance regression and temporal band-pass filtering (0.01–0.1 Hz) were performed. Nuisance regressors included detrending terms, motion parameters and their derivatives, tissue-derived signals, and selected non-brain PCA components, as detailed in the inset. The filtered data were subsequently normalized to template space and spatially smoothed using a 3-mm FWHM Gaussian kernel. DICOM: Digital Imaging and Communications in Medicine, NIfTI: Neuroimaging Informatics Technology Initiative, PCA: principal component analysis.

